# Mechanistic insights of glucosome condensate formation by stochastic modeling approaches

**DOI:** 10.1101/2022.06.27.497813

**Authors:** Hye-Won Kang, Luan Nguyen, Songon An, Minjoung Kyoung

**Affiliations:** Department of Mathematics and Statistics, University of Maryland Baltimore County (UMBC), 1000 Hilltop Circle, Baltimore, MD, 21250; Department of Chemistry and Biochemistry, University of Maryland Baltimore County (UMBC), 1000 Hilltop Circle, Baltimore, MD, 21250; Program in Oncology, Marlene and Stewart Greenebaum Comprehensive Cancer Center, University of Maryland, Baltimore, MD 21201

**Keywords:** molecular dynamics, LAMMPS, metabolic condensates, liquid-liquid phase separation, stochastic model, glucose metabolism, phosphofructokinase, glucosome

## Abstract

Human liver-type phosphofructokinase 1 (PFKL) has been shown to play a scaffolder role to recruit and organize glycolytic and gluconeogenic enzymes into a multienzyme metabolic condensate, the glucosome, that regulates glucose flux in living human cells. However, it has been challenging to characterize which factors control phase separation of PFKL and so glucosome condensates in a living cell, thus hampering to understand a mechanism of reversible glucosome formation and its functional contribution to human cells. In this work, we developed a stochastic model *in silico* using the principal of Langevin dynamics to investigate how biological properties of PFKL contribute to the formation of glucosome condensates. Molecular dynamics simulation using the Large-scale Atomic/Molecular Massively Parallel Simulator (LAMMPS) revealed the importance of an intermolecular interaction between PFKLs, an effective concentration of PFKL at a region of interest, and a pre-organization of its own self-assembly in formation of PFKL condensates and control of their sizes. Such biological properties that define intracellular dynamics of PFKL appear to be essential for phase separation of PFKL and thus formation of glucosome condensates. Collectively, our computational study provides mechanistic insights of glucosome formation, particularly an initiation step through the formation of PFKL condensates in living human cells.

## Introduction

Biomolecular condensates formed by liquid-liquid phase separation (LLPS) organize biomolecules such as proteins and/or nucleic acids spatially and temporally in cellular cytoplasm or nucleoplasm (Banani et al., 2017; Shin and Brangwynne, 2017). Their functional significances have been identified to include, but not limited to, regulation of specific biochemical reactions, sensitive sensing of cellular environmental changes, localization of specific molecules into defined subcellular locations, and/or buffering concentrations of biomolecules in cells (Alberti et al., 2019). In addition, pathological importance of biomolecular condensates have been recognized particularly in human neuronal diseases, for example, Alzheimer’s disease (Ambadipudi et al., 2017; Brundin et al., 2010), Parkinson’s disease (Shulman et al., 2011), and amyotrophic lateral sclerosis (Robberecht and Philips, 2013). Therefore, it is critical to understand molecular-level mechanisms of how the formation of biomolecular condensates are initiated and thus functionally regulated in living cells.

Of particular, intermolecular interactions between biomolecules and macromolecular crowding effects have been examined as examples of important factors that drive biomolecules to form self-assembled condensates (Banani *et al*., 2017; Fare et al., 2021; Hancock, 2004; Peran and Mittag, 2020; Schuster et al., 2021; Walter and Brooks, 1995). Thermodynamically, an intricate balance between biomolecule-biomolecule interactions and biomolecule-water interactions would determine the solubility of biomolecules in cells (Banani *et al*., 2017). Meanwhile, macromolecular crowding decreases an available volume to any biomolecules due to the volume occupancy by other macromolecules (i.e., crowders). In other words, crowders increase an effective concentration of biomolecules (Kuznetsova et al., 2014) and change the activity of water or valency of interacting biomolecular partners (Alberti *et al*., 2019), which influence on phase separation in cells. Nevertheless, until recently (Paloni et al., 2020; Sanders et al., 2020), it has remained largely elusive how these parameters and others that are mostly characterized by polymer chemistry or engineered model systems are related to various functional roles of biomolecular condensates in cells (Banani *et al*., 2017; Fare *et al*., 2021; Peran and Mittag, 2020; Schuster *et al*., 2021; Walter and Brooks, 1995).

At the same time, molecular dynamics (MD) simulation has been recently employed to understand LLPS-mediated formation of biomolecular condensates. Particularly, various properties that influence biomolecules and/or their surroundings, including for instance concentrations, multivalency, stoichiometry, temperature, and/or salt concentrations, have been evaluated *in silico* to understand their influence on phase behaviors of biomolecular condensates (Chou and Aksimentiev, 2020; Espinosa et al., 2020; Pyo et al., 2022; Ronceray et al., 2022; Tsanai et al., 2021; Zhang et al., 2021). Additionally, impacts of intrinsically disordered domains and nucleic acids have been simulated to understand their sequence-specific or structural roles on formation of biomolecular condensates (Abyzov et al., 2022; Alshareedah et al., 2021; Kaur et al., 2021; Lin et al., 2022; Mompean et al., 2021; Shillcock et al., 2020; Shillcock et al., 2022; Zheng et al., 2020). Therefore, MD simulations have been primarily employed to promote the influence of these properties on phase behavior of condensates or their formation rather than to decipher biological and physiological significance of these properties that would govern functional roles of condensates.

Enzymes in metabolic pathways have been demonstrated to form multienzyme metabolic condensates in living cells (Schmitt and An, 2017). Specifically, in human cells, enzymes that catalyze glucose metabolism (i.e., glycolysis and gluconeogenesis) and *de novo* purine biosynthesis, are spatially organized into multienzyme metabolic condensates, namely the glucosome (Kohnhorst et al., 2017) and the purinosome (An et al., 2008), respectively, to regulate desired metabolic flux at subcellular levels. Subsequent biophysical characterization has revealed that the pathway enzymes are dynamically partitioned into glucosomes or purinosomes and freely exchanged with surroundings, indicating no lipid membrane around the condensates (Kohnhorst *et al*., 2017; Kyoung et al., 2015). While glucosomes are characterized to form in various sizes and shapes (Kohnhorst *et al*., 2017), purinosomes are mostly organized spherically in one size category (An *et al*., 2008). Briefly, glucosomes are categorized into three subgroups; small-sized glucosomes are defined to have less than a calculated area of a point spread function for the emission of monomeric enhanced green fluorescent protein (i.e., ∼ 0.1 μm^2^), medium-sized glucosomes have between 0.1 μm^2^ and 3 μm^2^ since non-cancerous human cells do not display glucosomes in larger than 3 μm^2^ from our conditions, and thus large-sized glucosomes are defined to have larger than 3 μm^2^ that have been found in various cancer cells (Kohnhorst *et al*., 2017). Meanwhile, purinosomes generally display in similar sizes as medium-sized glucosomes in HeLa and Hs578T cells (Kohnhorst *et al*., 2017). Nevertheless, unlike other biomolecular condensates, it has been difficult to determine which LLPS-promoting biological and/or physiological factors would contribute to the formation of metabolically active glucosomes and purinosomes in human cells due to the lack of *in vitro* functional reconstitution strategies.

Mathematical modeling and *in silico* strategies have been initiated to understand mechanisms of glucosome formation in the cytoplasm of human cells (Davis et al., 2015; Jeon et al., 2018). Previously, we have developed a mathematical model to address glucosome formation in various sizes and their size-dependent functions using ordinary differential equations (Jeon *et al*., 2018). With the model, we have demonstrated how glucosome condensates regulate glucose flux between energy metabolism (i.e., glycolysis) and building block biosynthesis (i.e., the pentose phosphate pathway and serine biosynthesis) in a condensate size-dependent manner. However, the model does not address the importance of biological parameters of individual enzymes during the formation of glucosome condensates, which include, but not limited to, their diffusions, frictions, and interactions in a living cell. Therefore, no computational work has aimed to build a mathematical model to determine which and how biologically relevant parameters contribute to the formation of glucosomes and their functional roles.

In this work, we developed a stochastic model using the principle of Langevin dynamics to investigate the importance of intracellular properties of a scaffolder enzyme during the formation of glucosome condensates. We simulated our model using the Large-scale Atomic/Molecular Massively Parallel Simulator (LAMMPS) (Plimpton, 1995; Thompson et al., 2022), which embeds the parallel spatial-decomposition, neighbor-finding, and communication algorithms. Specifically, we examined the impacts of enzyme-enzyme interaction strengths, enzyme concentrations, alterations of enzyme properties, and a pre-organization of enzyme’s initial self-assembly on the formation of PFKL condensates. Consequently, we found that biological parameters that would dictate intracellular dynamics of PFKL are fine-tuned to promote phase separation of PFKL in cells. Considering that PFKL plays a scaffolder role during glucosome formation (Kennedy et al., 2022; Kohnhorst *et al*., 2017; Webb et al., 2017), our computational study reveals molecular-level mechanisms by which glucosome formation is initiated in living human cells.

## Methods

### Development of a stochastic model

Human PFKL catalyzing the conversion of fructose-6-phosphate to fructose-1,6-bisphosphate in glycolysis has been demonstrated to play a scaffolder role in formation of a multienzyme glucosome condensate (Kennedy *et al*., 2022; Kohnhorst *et al*., 2017). Our modeling studies focused on intracellular properties of PFKL and their roles in condensates formation. Since PFKL is catalytically active as a tetramer, we simplified a PFKL tetramer as a point particle in our model. In our model, we also set a region of interest in the cytoplasm as a cubic domain [−*L*/2,*L/*2]^3^. We defined a position vector and a velocity vector of the *i*th PFKL tetramer as ***r***_*i*_ and *ν*_*i*_, respectively, and denoted a mass of each PFKL tetramer as *m*, a friction constant as *γ*, and an isotropic diffusion coefficient as *D*.

Our stochastic model was built using the principle of Langevin dynamics with a pairwise interaction expressed by the Lennard-Jones potential (Erban and Chapman, 2019; Leimkuhler and Matthews, 2016). The position information of the *i*th PFKL tetramer was expressed as

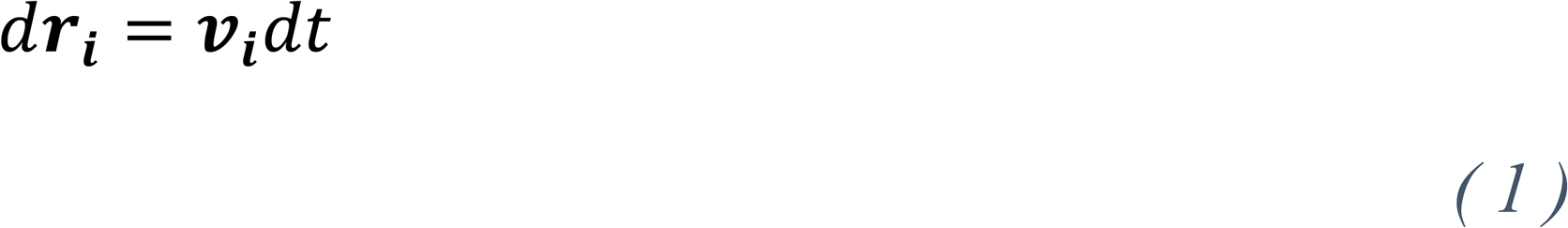

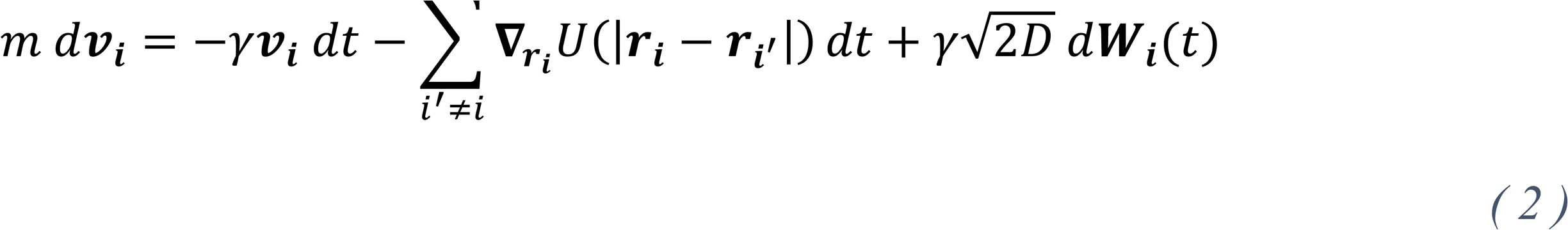

where ***W***_*i*_(*t*) is a standard Brownian motion in three dimensional space. Three terms on the right-hand side of Eq. (2) represent a frictional force, a pairwise enzyme-enzyme interaction force, and a random force applied to the *i*th PFKL tetramer. We chose a periodic boundary condition to describe a subcellular region of interest as an open system. The Lennard-Jones potential that describes the interaction between each pair of PFKL tetramers was expressed as

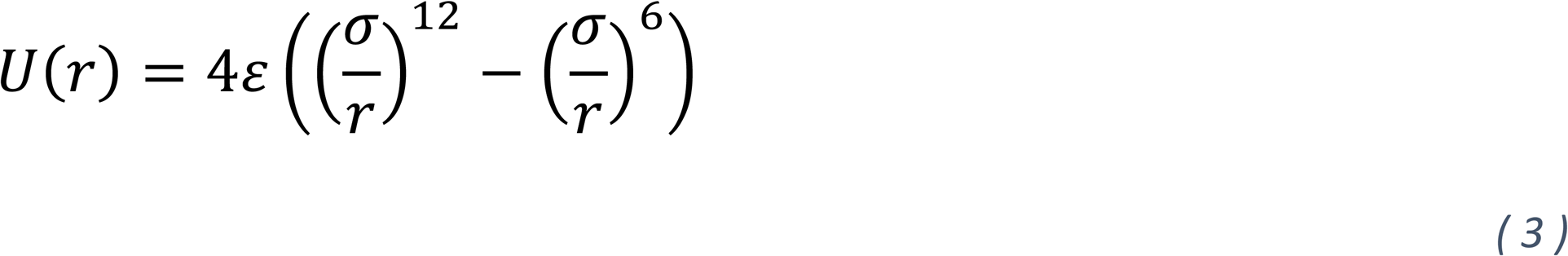

where *r* represents the distance between a pair of PFKL tetramers when they interact with each other. Accordingly, the depth of the potential well (*ε*) indicates how strongly two PFKL tetramers interact, and σ is a distance at which the potential energy *U* becomes zero.

In the LAMMPS, the Langevin dynamics is governed by

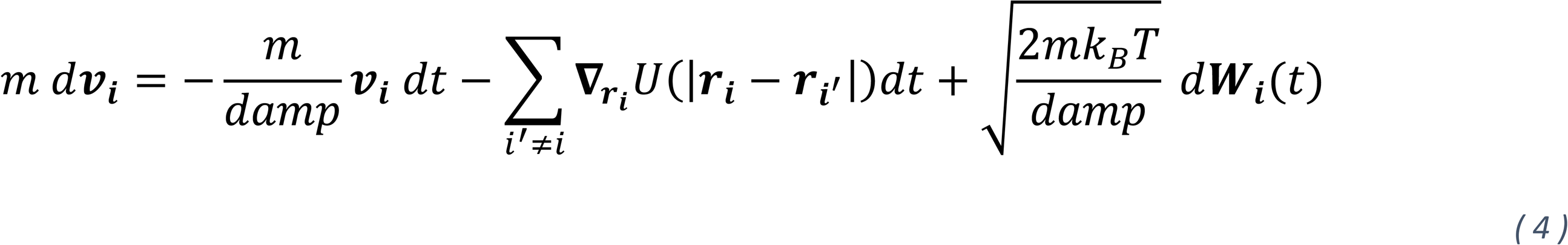

where *k*_*B*_ is the Boltzmann constant and *T* is temperature. When Eq. (4) was compared with Eq. (2), a friction constant (*γ*) and a diffusion coefficient (*D*) were expressed in Eq. (4) as *γ = m/damp* and *D =* (*k*_*B*_*T*)*ddddmmdd/mm = k*_*B*_*T/γ*, respectively.

Then, to drive a dimensionless equation, we further defined scaled variables for time, position, velocity, and random movement as

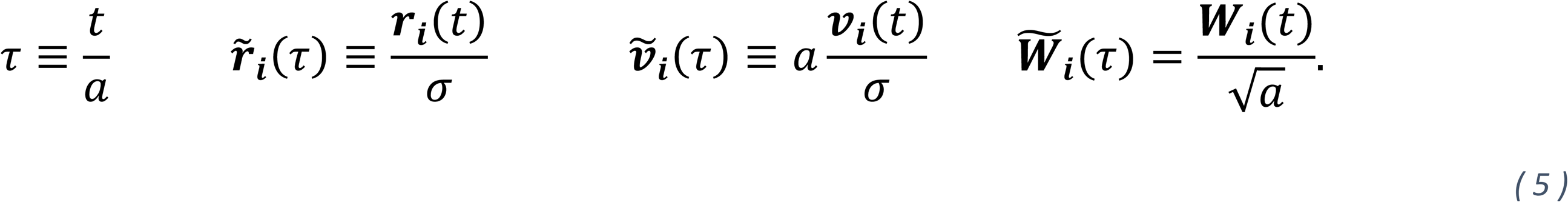

Plugging Eq. (3) in Eq. (4) and replacing *t* and *dt* in Eq. (4) by *a* τ *τ* and *adτ*, we obtained

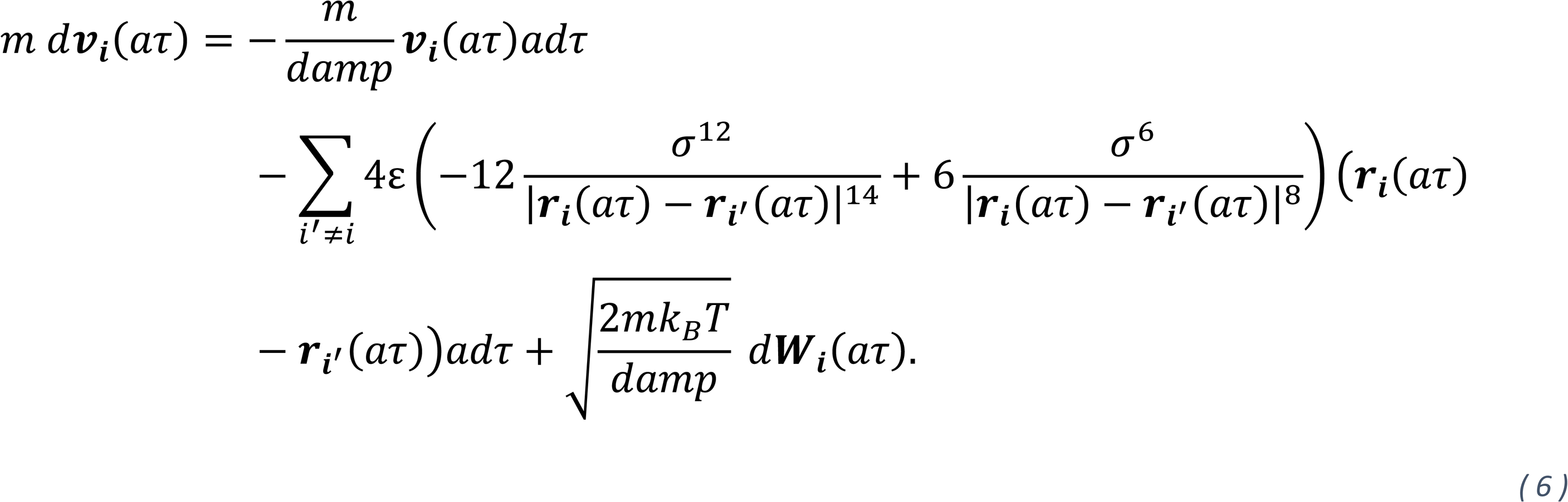

Using the scaled variables in Eq. (5), Eq. (6) became

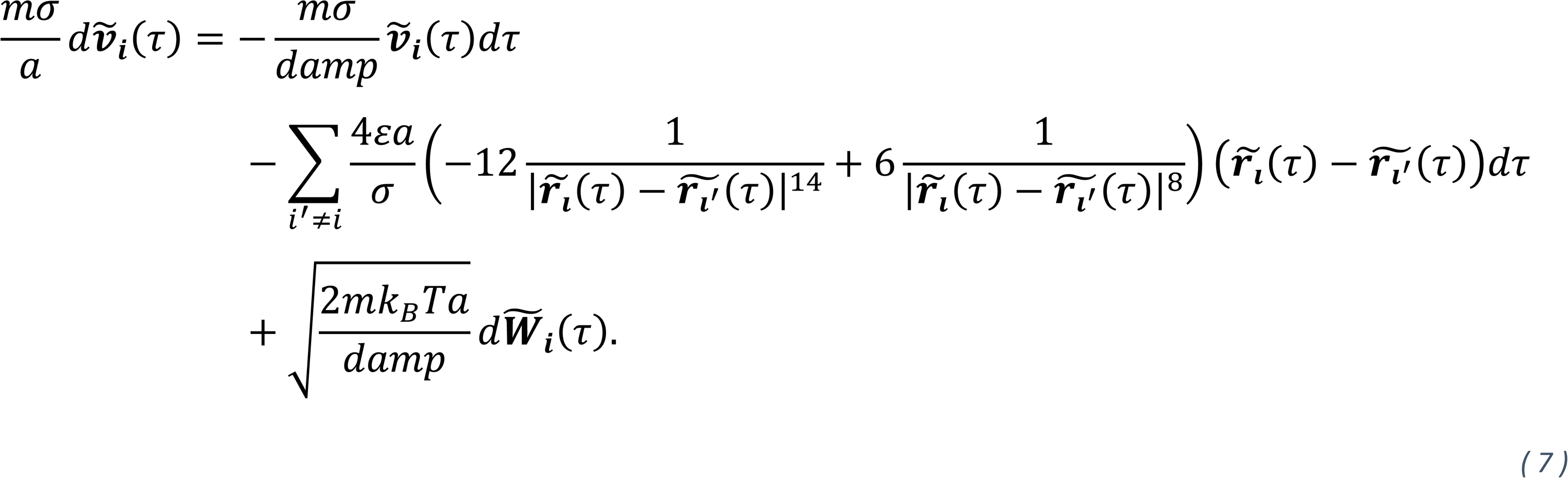

After defining 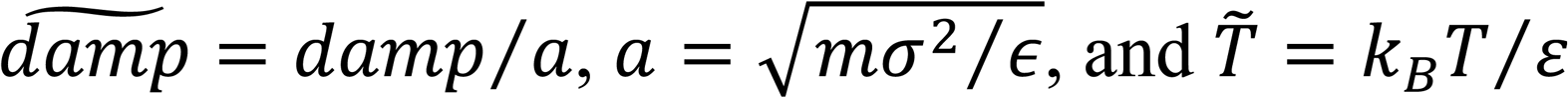, we obtained a scaled equation as

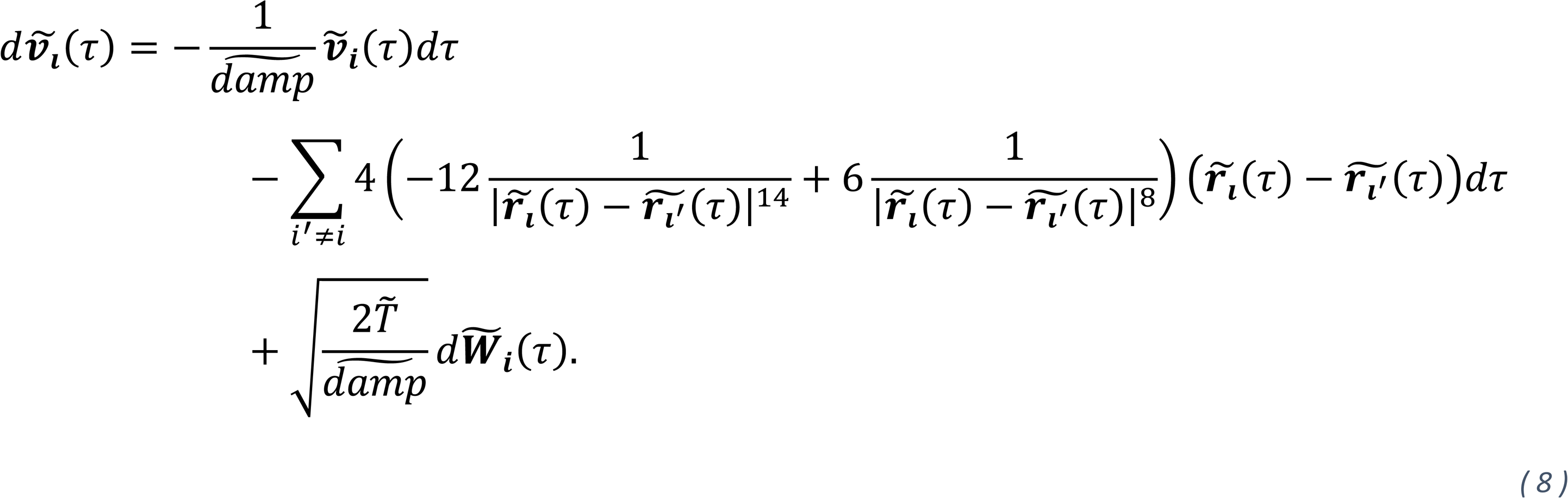

All simulations were then performed using the scaled equation (Eq. (8)) with parameters of 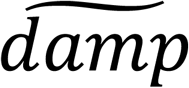, 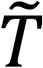, and *a*. Note that the LAMMPS codes were modified to include both multiple particles and polymers to accommodate coexistence of multiple PFKL tetramers and PFKL filaments in our simulation (Brackley, 2019).

### Parameter estimation of the model

We then determined various parameters for our model (**Table 1**). The intermolecular distance (*σ*) between a pair of PFKL tetramers was estimated as following. The dimension of a PFKL tetramer was obtained from X-ray crystal structure of human platelet-type PFK (PFKP) tetramer (Webb et al., 2015); 13.8 *nm* × 10.3 *nm* × 15.9 *nm*. PFKL tetramers were also anticipated to be spaced closely each other but still apart by a positive distance within PFKL condensates. We determined from intracellular fluorescence resonance energy transfer (FRET) experiments that the surface-to-surface distance between two PFKL tetramers was on average 5.5 *nm* inside glucosome condensates (Kennedy *et al*., 2022). Accordingly, we set the distance *σ*, 21.4 *nm*, which represents a center-to-center distance between two PFKL tetramer, including the FRET distance, at which the potential energy become zero. Next, based on the sequence of amino acids, the molecular weight of human PFKL tetramer was calculated as 718.84 *kDd*, thus determining the mass of each PFKL tetramer, *m*. A diffusion coefficient (*D*) of human PFKL, when they did not form PFKL condensates, was experimentally measured previously as 0.126 *±* 0.050 *μm*^2^*sec*^−1^ (Kohnhorst *et al*., 2017), establishing the parameter value of *D* in Eq. (2).

**Table 1.** Parameter values in the stochastic model for PFKL tetramers.

| Symbol | Names | Values |
| --- | --- | --- |
| $L$ | The edge of a cubic domain | $1.2849 \mu m$ |
| $N$ | A total number of PFKL tetramers | 512 |
| $\varepsilon$ | The potential well depth<br>(a.k.a. the intermolecular interaction strength) | $4.3 \times 10^{-23} \sim 5.2 \times 10^{-23} kg \mu m^2 sec^{-2}$ |
| $\sigma$ | The distance between two particles when<br>$U = 0$ | $0.021415 \mu m$ |
| $m$ | The mass of the PFKL tetramer | $5.64783 \times 10^{-22} kg$ |
| $\gamma$ | The friction coefficient of the PFKL tetramer | $1.4941 \times 10^{-22} kg sec^{-1}$ |
| $D$ | The diffusion coefficient of the PFKL tetramer | $0.126 \mu m^2 sec^{-1}$ |
| $t_{end}$ | End of the simulation time | 1800 sec |
| $\Delta t$ | Unit of each time step in the simulation | 0.001 sec |

Human PFKLs are also known to form glucosome condensates in various sizes by recruiting other glycolytic and gluconeogenic enzymes (Kohnhorst *et al*., 2017). Since small-or medium-sized glucosomes were defined to have less than 0.1 *μm*^2^or between 0.1*∼*3 *μm*^2^, respectively (Kohnhorst *et al*., 2017), the cubic domain size was determined to be large enough to contain PFKL condensates in small or medium sizes, accordingly setting a cubic domain [−*L/2,L/*2]^3^ where *L =* 60σ. We also decided to simulate 512PFKL tetramers (*N =* 512) that would occupy more than 10% of a total volume of the cubic domain. Each PFKL tetramer was then located at a cubic grid so that all PFKL tetramers were initially distributed uniformly in the cubic domain. Then, when more than 9 PFKL tetramers were assembled during our MD simulation, we counted it as a condensate because 3-8 PFKL tetramers were shown to form filaments *in vitro* (Webb *et al*., 2017). Next, to determine our simulation time, we considered two facts; (i) that LLPS has been reported to occur in the time scale of several seconds to minutes (Dundr et al., 2004; Phair and Misteli, 2000; Weidtkamp-Peters et al., 2008) and also (ii) that PFKL condensates have been experimentally shown to form within minutes to several hours after addition of exogenous cues (Jeon et al., 2022a; Jeon *et al*., 2018; Jeon et al., 2022b; Kennedy *et al*., 2022; Kohnhorst *et al*., 2017). After careful evaluation of various simulation times ranging from 15 to 60 minutes, we decided to run our MD simulations for 30 minutes because it allowed us to robustly evaluate whether a given set of input parameters would promote the formation of PFKL condensates or not. Accordingly, we performed all simulations up to 30 minutes (*= t*_*end*_) with time step Δ*t* (Table 1), and then recorded the frequency (%) of PFKL condensates formation and, when they formed, their sizes during and after each simulation.

To simulate Eq. (1)-Eq. (3), we input additional parameters for a potential well depth (*ε*) and a friction coefficient (*γ*). First, we tested a wide range of potential well depth values *ε* to discover the threshold where the phase separation of PFKL occurred. Afterwards, we simulated with the neighboring parameter values to investigate how PFKL condensates formation was affected by. Second, to estimate a friction coefficient, we surveyed a range of propulsion velocity of various biomolecules as in the order of 10*∼*10^3^*μmsec*^−1^ (Feng and Gilson, 2019; Jee et al., 2018; Santiago and Simmel, 2019). At the same time, some enzymes appear to show moderate to least amount of self-propulsion so that they were observed to move with a speed in a range of 0.1*∼*2.5 *μmsec*^−1^ (Arque et al., 2019). When we then assumed that each PFKL tetramer was only governed by a frictional force and a random force, the mean squared velocity of each PFKL tetramer was expressed as 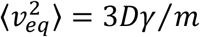 (Medved et al., 2020) with a value of 0.1 *μm*^2^*sec*^−2^. Accordingly, the friction coefficient *γ* was calculated as 1.4941 × 10^−22^*kgsec*^−1^ (Table 1). Note that the chosen friction coefficient provides a velocity relaxation time as *m/γ ≈* 4 *sec* when we ignore an enzyme-enzyme interaction force applied to each PFKL tetramer.

## Results

Biomolecular condensates driven by LLPS in living systems often require adequate interactions between biomolecules, subcellularly concentrated biomolecules, and multivalent biomolecular interactions to maintain their dynamic properties. In this work, we developed and simulated a stochastic model *in silico* using the LAMMPS (Plimpton, 1995; Thompson *et al*., 2022)to understand the role of each biological parameter that the scaffolder enzyme of the glucosome (i.e., PFKL) would provide during the formation of glucosome condensates. Accordingly, we considered various scenarios that took into accounts of an intermolecular strength between PFKLs, an effective concentration of PFKL in a region of interest, an existence of second PFKL species due to alterations of PFKL properties at given environment, and the shape and flexibility of multivalent PFKL oligomers. Then, we analyzed two outputs in details from our MD simulation: the formation frequency of PFKL condensates and, when they formed, their size distribution. Please note that PFKL was shown to form filaments, at least *in vitro*, at certain conditions and ∼85 % of such filaments were composed by 3-8 PFKL tetramers *in vitro* (Webb *et al*., 2017). Accordingly, when more than nine PFKL tetramers were assembled during our MD simulation, we defined it as a PFKL condensate. Likewise, the largest condensate in size in our simulation would be a condensate that would have 512 PFKL tetramers. At the same time, understanding size distributions of PFKL condensates is important because the size of a PFKL condensate represents its functional contribution to glucose metabolism in human cells (Jeon *et al*., 2018; Kohnhorst *et al*., 2017). Collectively, our work here would provide mechanistic insights of how PFKL and thus glucosomes would form in various sizes to regulate glucose flux in human cells.

### Impacts of intermolecular interaction strengths on the formation of PFKL condensates

We first investigated how different strengths of the PFKL-PFKL interaction affect the formation of PFKL condensates. To begin with, all 512PFKL tetramers were considered indistinguishable from each other, indicating that their mass, frictional force, interaction strength, and diffusion coefficients are identical. We then gradually increased intermolecular interaction strengths between each pair of PFKL tetramers by changing *ε* values from 4.3 × 10^−23^ to 5.2× 10^−23^. The range of *ε* was determined to not only observe phase separation of PFKL but also allow us to quantify the formation of PFKL condensates in our MD simulation. The formation frequency of PFKL condensates was calculated by counting the number of occurrences that PFKL condensates appeared out of 300 independent runs of our simulations at each value of *ε* (**Figure 1A**). The sizes of PFKL condensates were also obtained by counting the number of PFKL tetramers forming each condensate at the end of the simulation time (i.e., 1800 *sec*) (**Figure 1B**).

**Figure 1.**
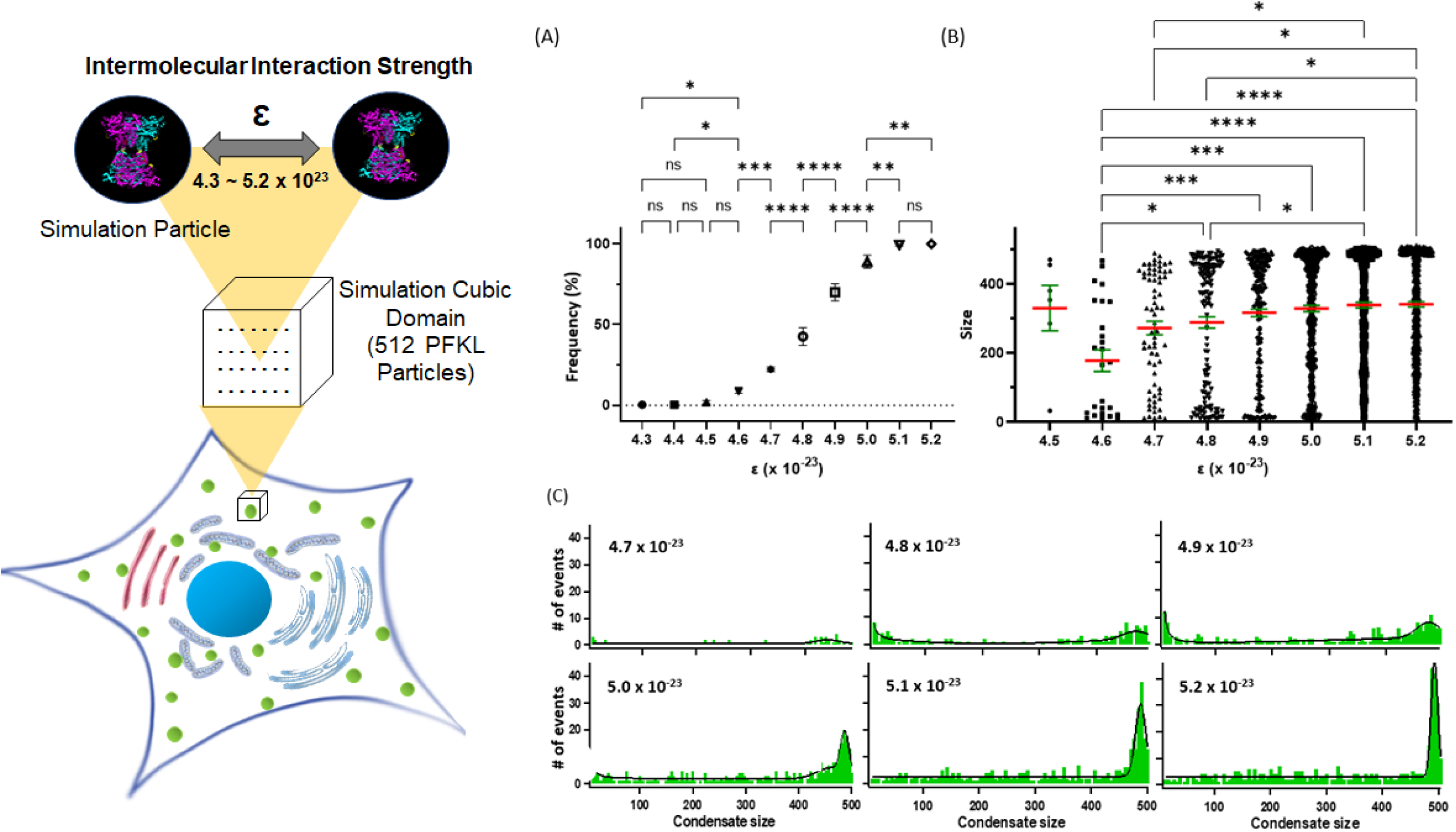
Impacts of intermolecular interaction strengths on the formation of PFKL condensates. While we gradually increased enzyme-enzyme interaction strengths from *ε* = 4.3 × 10^−23^ to *ε* = 5.2× 10^−23^, the formation frequency and the size distribution of PFKL condensates were measured and analyzed. (A) The frequencies of condensates formation and their standard errors at each interaction strength were plotted from 300 independent runs of simulations. (B) The size distributions of PFKL condensates were also shown at each interaction strength. Mean sizes of condensates (red lines) and standard errors of their sizes (green lines) at each interaction strength were graphed from 300 independent runs of simulations. (C) The size distributions of PFKL condensates at each interaction strength was also fitted using multi-modal functions (i.e., double gaussian and exponential). The number of PFKL condensates formed at a given size is graphed as green bars and the fitted size distributions are shown by black curves. Statistical analyses were performed using a two-way analysis of variance (ANOVA) along with Tukey’s multiple comparison tests. Statistical significance is defined as *p* < 0.05 with a 95% confidence interval while ‘ns’ refers to not significant. * *p* < 0.05, ** *p* < 0.01, *** *p* < 0.001, **** *p* < 0.0001.

As shown in **Figure 1A**, the stronger enzyme-enzyme interaction was simulated, the more frequent formation of PFKL condensates was observed. Briefly, when *ε* = 4.3 × 10^−23^ and *ε* = × 10^−23^, PFKL condensates were rarely produced (*<* 1%). However, at *ε* = 5.1 × 10^−23^ and *ε* = 5.2× 10^−23^ PFKL tetramers were assembled into condensates in most runs of simulations (*>* 99%). Otherwise, we detected coexistence of individual PFKL tetramers and PFKL condensate(s) at the end of simulations. In addition, we compared sizes of PFKL condensates with a range of *ε* from 4.5 × 10^−23^ to 5.2× 10^−23^ (**Figure 1B**). We found that the mean size of PFKL condensates increased as the interaction strength enhanced (red lines, **Figure 1B**). Note that in the case of *ε* = × 10^−23^, the mean size of PFKL condensates appeared to be biased because only 6 out of 300 runs of simulation resulted in the formation of PFKL condensates. Nevertheless, since the size distributions of PFKL condensates appeared to be too broad at each *ε* value, we further analyzed the data by fitting with multi-modal functions (i.e., double gaussian and exponential) (black lines, **Figure 1C**). Then, we noticed that as the interaction strength between PFKLs (*ε*) increased from × 10^−23^ to 5.2× 10^−23^, the mean size of a subpopulation of PFKL condensates that contained 400 to 512 PFKL tetramers increased from 441.30 ± 5.42 to 495.73 ± 1.18 and the width of the subpopulation’s distribution became narrower from 32.49 ± 8.39 to 8.49 ± 1.18 (**Figure 1B and 1C**). This result indicates that, as the interaction strength is increased within the range of our interest, condensates having 400 to 512 of PFKL tetramers appear to be rapidly promoted by condensational growth that would narrow the size distribution, rather than collisional growth that would increase the width of condensates’ size distribution (Lamb and Verlinde, 2011; Preuppacher and Klett, 2012). Meanwhile, we also noticed that a proportion of the subpopulation of small-sized condensates that contained less than 100 PFKL tetramers was reduced when the interaction strength between PFKLs was increased from *ε* = 4.6 × 10^−23^ to *ε* = 5.2× 10^−23^ (**Figure 1B**). Therefore, our MD simulation supports that the stronger enzyme-enzyme interaction is presented, the more frequent formation and the larger sizes of enzyme condensates are being formed.

### Correlation between an effective concentration of PFKL and its condensate formation

We then investigated the effect of a concentration of PFKL tetramers at a region of interest. In this work, we started our simulation with undistinguishable 512 PFKL tetramers in the cubic domain. However, to increase an effective concentration of PFKL, we decreased the volume of the simulation cubic domain by every 10% from 100% (i.e., 1x) to 60% (i.e., 0.6x) of its default size. We simulated this case at two different interaction strengths of *ε* = 4.5 × 10^−23^ (red in **Figure 2**) and *ε* = 4.7 × 10^−23^ (green in **Figure 2**) because at those *ε* values the frequencies of PFKL condensates formation were relatively low from the default size of our simulation domain (as shown in **Figure 1A**) and thus the concentration effect would be meaningfully detected in our simulation. Indeed, we observed at both scenarios that the formation frequencies of PFKL condensates reached ∼100% from only ∼2% at *ε* = 4.5 × 10^−23^ (i.e., 6 cases out of 300 simulations) or ∼22% at *ε* = 4.7 × 10^−23^ (i.e., 67 cases out of 300 simulations) (**Figure 2A**). We then compared condensate size distributions at different simulation volumes. Our data revealed that decreasing the simulation volume drastically increased the mean size of PFKL condensates (**Figure 2B**), indicating that the higher effective concentration of PFKL at a region of interest would be, the larger size of a PFKL condensate would be formed. Therefore, our results reveal a positive correlation between an effective concentration of an enzyme and its tendency to form a condensate as well as a physical size of its condensate.

**Figure 2.**
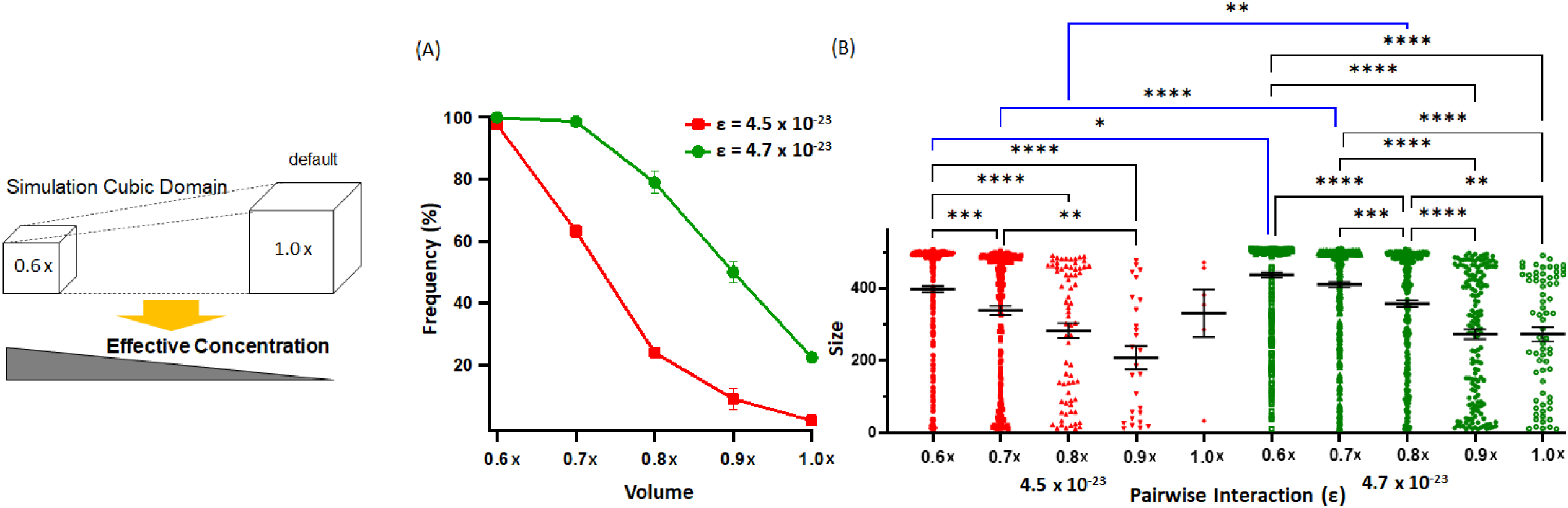
Correlation between an effective concentration of PFKL and its condensate formation. To simulate effects of various concentrations of PFKL at a given region of interest, we gradually reduced a simulation domain size from its default (1.0 ×) to its 60% volume (0.6 ×). The frequency of PFKL condensates formation (A) and their size distribution (B) were measured and analyzed at each simulation domain. Mean sizes of condensates (black lines) and standard errors (black error bars) are from 300 independent runs of simulations at each domain volume for two intermolecular interaction strengths of *ε* = 4.5 × 10^−23^ (red) and *ε* = 4.7 × 10^−23^ (green). Statistical analyses were performed using a two-way ANOVA along with Tukey’s multiple comparison tests. Statistical significance is defined as *p* < 0.05 with a 95% confidence interval. * *p* < 0.05, ** *p* < 0.01, *** *p* < 0.001, **** *p* < 0.0001.

### Potential effects of existence of second PFKL species as being part of a pre-organization

Meanwhile, metabolic enzymes have been discovered to form various spatially resolved structures in living cells. In facts, mesoscale filaments, rods, rings, sheets, lattices, tubes as well as liquid-like condensates have been observed under fluorescence microscopy and/or electron microscopy (Pilhofer and Jensen, 2013; Wilson and Gitai, 2013). Such mesoscale structures formed by metabolic enzymes may be constructed from nanoscale pre-organizations. Accordingly, we hypothesized that, if PFKL was organized into any form of pre-organization, each PFKL tetramer unit in a pre-organization might be regulated to present a stronger enzyme-enzyme interaction or a heavier mass than ones that freely diffusing PFKL tetramers would do, thereby resulting in the formation of PFKL condensates. Thus, we evaluated the effects of potential existence of second PFKL species in our simulations.

First, we assumed that two different species of PFKL tetramers would possess different levels of an intermolecular interaction strength (**Figure 3A**). In this simulation, we assigned 128 PFKL tetramers out of 512 PFKLs as the second species (i.e., species 2) to have an ability to provide a stronger enzyme-enzyme interaction than the interaction strength the rest of PFKL tetramers have (i.e., 384 PFKLs, species 1). Mathematically, we denoted the enzyme-enzyme interaction between species *i* and species *j* as *ε*_*ij*_. Then, we considered three different scenarios, including a control, with various enzyme-enzyme interaction levels. First, we took account of a control situation, as shown in **Figure 1**, where PFKL tetramers in pre-organization and individual PFKL tetramers had the same enzyme-enzyme interaction strength. Relative levels of the interaction strengths between species *i* and species *j* were then expressed as *ε*_11_: *ε*_12_: *ε*_22_*=* 1: 1: 1 where *ε*_*ij*_ *= ε*_0_, *i, j =* 1,2. Next, we mimicked a situation that PFKL tetramers in pre-organization were attracted to each other with a 20% stronger enzyme-enzyme interaction than a default interaction strength between individual PFKL tetramers. In this situation, we further assumed an interaction level between each PFKL tetramer in pre-organization and an individual PFKL tetramer to be 10% different or no different from the default interaction strength, resulting in simulation sets of *ε*_11_: *ε*_12_: *ε*_22_=1:1.1:1.2 or *ε*_11_: *ε*_12_: *ε*_22_*=* 1: 1: 1.2, respectively. Note that 20% additional strength was assigned between species 2 PFKLs because it was mathematically enough to see a significant increase in the frequency of PFKL condensates formation. Simulating these cases at three different default interaction strengths (i.e., *ε*_0_ *=* 4.6 × 10^−23^, 4.7 × 10^−23^, and × 10^−23^) made it clear that the impact of the existence of species 2 was indeed significant on the formation of PFKL condensates (**Figure 3A**). We conclude that species 2-containing pre-organization may be able to initiate to form small condensates, which would help in turn to form larger PFKL condensates.

**Figure 3.**
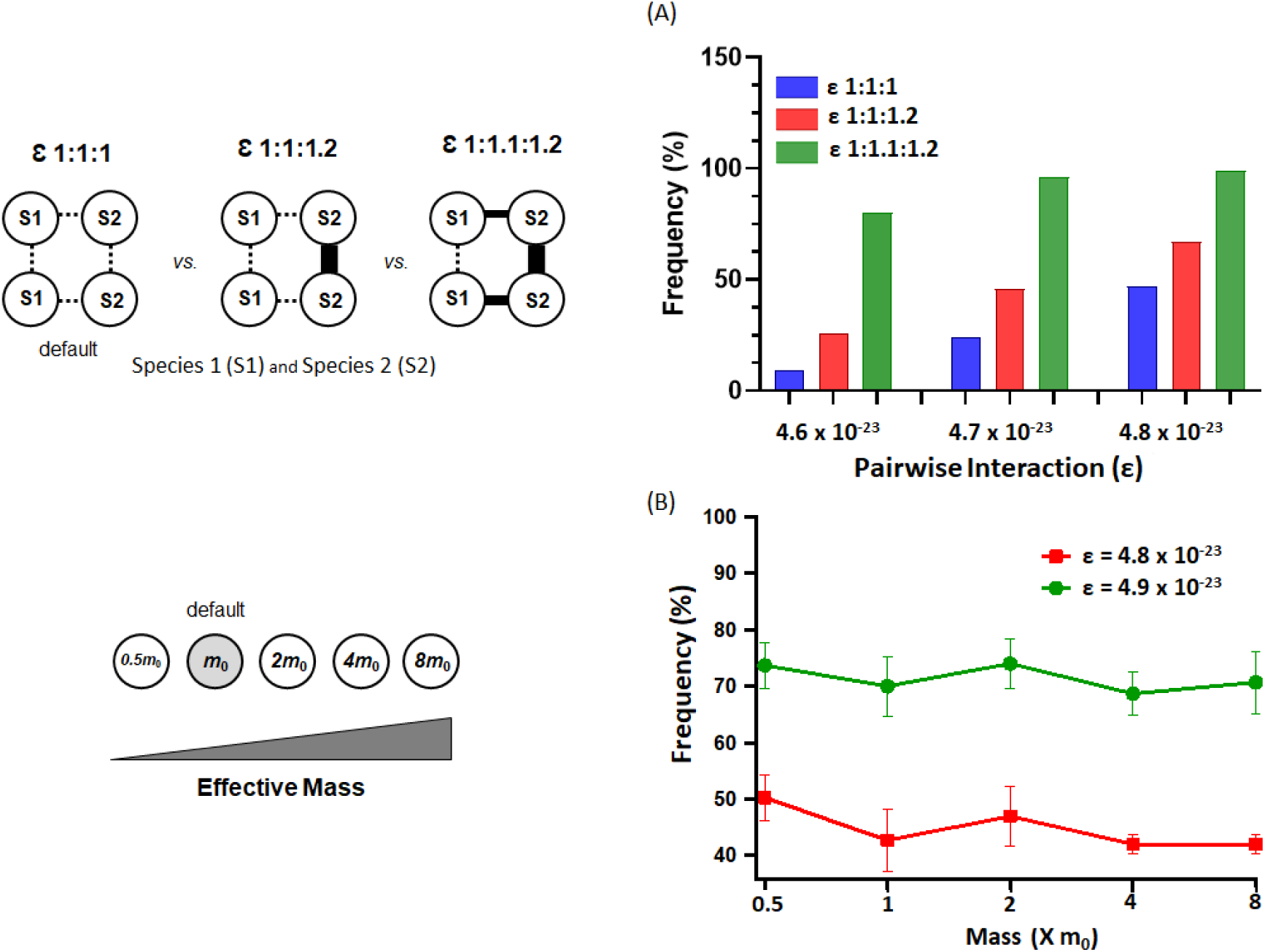
Potential effects of existence of second PFKL species as being part of a pre-organization. We assumed that if PFKL tetramers would form pre-organization prior to condensates formation, there could be at least two species of PFKL tetramers where one might have a stronger interaction strength or display a heavier mass than the other might have. (A) First, we expressed the enzyme-enzyme interaction strength between species *i* and species *j* as *ε*_*ij*_, *i, j =* 1,2. Then, three scenarios were considered when ratios of intermolecular interaction strengths between two species of PFKL were *ε*_11_: *ε*_12_: *ε*_22_=1:1:1 (blue), *ε*_11_: *ε*_12_: *ε*_22_=1:1:1.2 (red), or *ε*_11_: *ε*_12_: *ε*_22_=1:1.1:1.2 (green). Each scenario was tested in the case of species 1 possessing an enzyme-enzyme interaction strength of *ε*_11_ *=* 4.6 × 10^−23^, 4.7 × 10^−23^, or 4.8 × 10^−23^. The frequency of PFKL condensates formation is from 100 independent runs of simulations for each case. (B) Second, we simulated a case in which two species of PFKL tetramers might contribute different masses to condensate formation. Species 1 had a default value of the mass (*m*0), and thus species 2 had 0.5, 1, 2, 4, or 8 times of the default value of the mass. Both species had the same level of an enzyme-enzyme interaction strength, denoted as *ε*. The frequencies of PFKL condensates formation and their standard errors were from 300 independent runs of simulations for each case at *ε* = 4.8 × 10^−23^ (red) or *ε* = 4.9 × 10^−23^ (green).

Second, we introduced a heavier mass species to distinguish PFKL tetramers in pre-organization from individual PFKL tetramers. In this simulation, we examined whether different diffusions due to different masses of PFKL tetramers would affect the formation of PFKL condensates. Similarly, we considered 384 PFKL tetramers having a default mass (*m*_0_) and 128 PFKL tetramers with a mass that was equal to 0.5*m*_0_, *m*_0_, 2*m*_0_, 4*m*_0_, or 8*m*_0_. Other than the mass, both species of PFKL tetramers were assumed to have the same physical properties. We then evaluated each scenario when the enzyme-enzyme interaction strength was equal to *ε* = 4.8 × 10^−23^ or *ε* = 4.9 × 10^−23^. Note that the two *ε* values were chosen so that individual PFKL tetramers and PFKL condensates would coexist during simulations when all PFKL tetramers were assumed to have a default mass *m*_0_ as a control scenario shown in **Figure 1**. When two species of PFKL tetramers had the same default mass (*m*_0_), the frequency of PFKL condensates formation were 42.7% and 70.0% at *ε* = 4.8 × 10^−23^ and *ε* = 4.9 × 10^−23^, respectively (**Figures 1A** and **3B**). However, as we varied the mass of species 2 PFKL tetramer, we did not see a significant change in the frequency of PFKL condensates formation (**Figure 3B**). Therefore, we conclude that the formation of PFKL condensates may not significantly depend on additional PFKL species displaying different masses.

### Significant and tunable contribution of a defined pre-organization of PFKL to the formation of condensates and their sizes

Nevertheless, multivalent polymers such as filamentous compounds have been known to self-organize into higher-ordered structures (Banani *et al*., 2017; Harmon et al., 2017). At the same time, PFKL tetramers have been previously demonstrated to form nanoscale filaments *in vitro* (Foe and Trujillo, 1980; Webb *et al*., 2017) and have four contact sites per tetramer while being in a filament (Webb *et al*., 2017). With a possibility that PFKL filaments might act as multivalent polymers, we hypothesized that PFKL filaments would have more contact sites (i.e., higher multivalency) than an individual PFKL tetramer, thus increasing the probability of PFKL condensates formation. To test the hypothesis, we introduced two variables, the flexibility and the length of PFKL filaments, in our MD simulations.

We first defined that species 1 was an individual PFKL tetramer and species 2 was an oligomeric PFKL filament (**Figure 4**). A total of four PFKL filaments were used in each simulation, where each PFKL filament was composed of multiple numbers of PFKL tetramers (i.e., 3, 4, 5, or 6), defining the length of each PFKL filament in our simulation. In all scenarios, the total number of PFKL tetramers regardless of their participation in filaments were maintained as 512. We then implemented different lengths of four PFKL filaments in two ways. One way was to model ‘elastic’ PFKL filaments as PFKL tetramers were connected via freely-rotating interaction points. The other way was to model ‘rigid’ PFKL filaments as PFKL tetramers were connected through fixed angles to maintain 180°. Additionally, our simulations at each scenario were performed at two different interaction strengths: *ε* = 3.3 × 10^−23^ and *ε* = 4.5 × 10^−23^. Note that we chose these two *ε* values so that no PFKL condensate formed in most runs with the shortest PFKL filaments containing 3 PFKL tetramer in one condition (i.e., *ε* = 3.3 × 10^−23^) while PFKL condensates were ensured to form (*ε* = 4.5 × 10^−23^).

**Figure 4.**
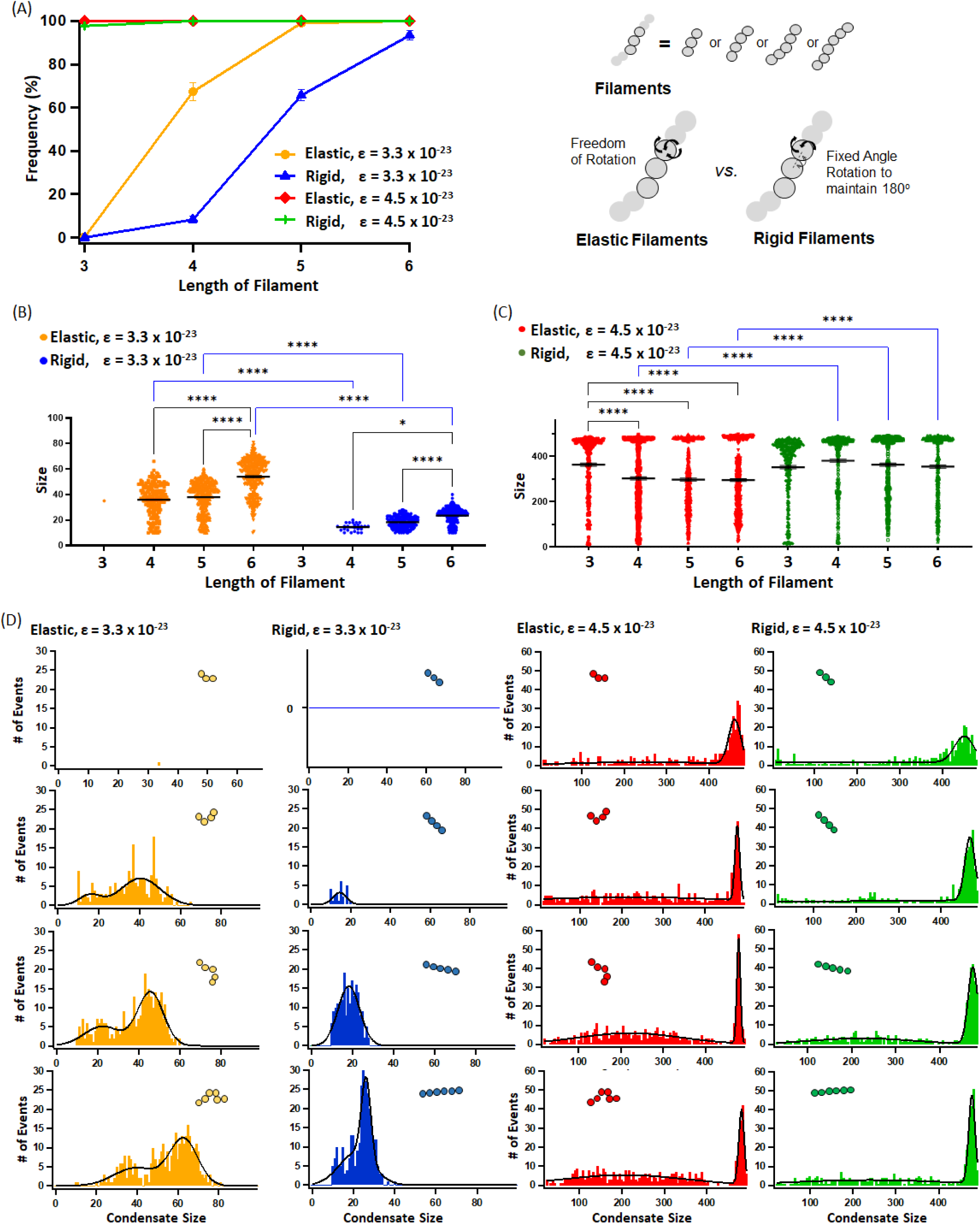
Contribution of a defined pre-organization of PFKL to the formation of condensates and their sizes. In this simulation, we assumed two species of PFKLs: species 1 was a freely diffusing individual PFKL tetramer whereas species 2 was an oligomerized PFKL filament consisting of 3 to 6 PFKL tetramers. Elastic (orange and red) or rigid (blue and green) filaments having different lengths were simulated at the enzyme-enzyme interactions of *ε* = 3.3 × 10^−23^ (orange and blue) or *ε* = 4.5 × 10^−23^ (red and green). 300 independent runs of simulations were performed at each case. (A) The frequency (%) of PFKL condensates formation was graphed as varying the flexibility and length of filaments. (B and C) The size distributions of observed PFKL condensates were displayed at each enzyme-enzyme interaction strength for each filament length. (D) The size distributions of PFKL condensates were also fitted using multi-modal functions. The numbers of PFKL condensates formed at a given size are graphed as bars and the fitted size distributions are shown by black curves. Statistical analyses were performed using a two-way ANOVA along with Tukey’s multiple comparison tests. Statistical significance is defined as *p* < 0.05 with a 95% confidence interval. * *p* < 0.05, **** *p* < 0.0001.

When we then performed 300 runs of each simulation, we observed positive correlations between the length of PFKL filaments and the formation frequency of PFKL condensates (**Figure 4A**). When the enzyme-enzyme interaction was set as *ε* = 4.5 × 10^−23^ at which PFKL tetramers were barely organized into PFKL condensates in the absence of a filamentous structure (**Figure 1**), it was clear that having a filamentous structure, regardless of its flexibility, made PFKL tetramers effectively organized into their condensates (red and green, **Figure 4A**). On the other hand, when the enzyme-enzyme interaction was set as very weak (i.e., *ε* = 3.3 × 10^−23^) at which condensates had no chance to be formed without a filamentous structure, having elastic PFKL filaments showed more beneficial to recruit individual PFKL tetramers than having rigid PFKL filaments (orange vs. blue, **Figure 4A**). Nevertheless, having high-ordered filament structures enhanced the probability of PFKL tetramers to form condensates. Overall, this result supports that filament-participating PFKL tetramers have thermodynamic advantages at the beginning of simulations and thus strengthen attractive forces by increasing its effective multivalency for freely diffusing PFKL tetramers during simulations.

Next, we analyzed size distributions in each case. When the enzyme-enzyme interaction was very weak (i.e., *ε* = 3.3 × 10^−23^) (**Figure 4B**), the sizes of PFKL condensates were overall larger with elastic filaments than the sizes with rigid filaments (orange vs blue, **Figure 4B**). This may be explained by our observation that during MD simulations, oligomerized PFKL tetramers in each elastic filament got more densely packed at the beginning of simulation due to their freedom of rotation than the ones in rigid filaments. Consequently, elastic filaments might recruit individual PFKL tetramers more effectively than rigid ones. In addition, when the intermolecular interaction between PFKLs is very weak (i.e., transient), regardless of the flexibility of filamentous structures, the length of PFKL filaments would be a determinant for condensate size.

However, we observed different trends when the enzyme-enzyme interaction was set as *ε* = 4.5 × 10^−23^ (**Figure 4C**). Condensates formed from elastic PFKL filaments were smaller in size on average than those from rigid PFKL filaments particularly when the filaments were formed by more than four PFKL tetramers (**Figure 4C**). This may be explained by our observation that each elastic filament started to recruit individual PFKL tetramers from the beginning of simulations and ended up forming multiple small condensates whereas rigid filaments initially got attracted by themselves and then recruited individual PFKL tetramers to form relatively larger condensates. Indeed, the number of PFKL condensates produced by elastic over rigid PFKL filaments was increased 16∼29%. Although we cannot rule out a possibility that multiple small PFKL condensates may merge into larger ones as time goes, it is important to note here that our simulation was performed consistently throughout our investigation with a fixed timescale and a fixed total number of PFKL tetramers. Moreover, the geometry of rigid filaments appears to offer more restricted environment for freely diffusing PFKL tetramers to be bound than the geometry of elastic filaments. Collectively, when the intermolecular interaction between PFKLs becomes relatively stable, the average sizes of PFKL condensates made by elastic PFKL filaments were smaller than the condensate sizes that were made by rigid filaments.

Additionally, we analyzed the size distributions of detected PFKL condensates in detail by fitting our data with multi-modal functions (i.e., double gaussian and exponential) (**Figure 4D**). First, when the enzyme-enzyme interaction was at *ε* = 3.3 × 10^−23^, the size distributions of PFKL condensates depended on the flexibility of the filaments (orange and blue, **Figure 4D**). Briefly, condensates formed from elastic filaments displayed two populations where the first population containing relatively small condensates was close to the size that could be formed by four filaments themselves being combined together while the second population was composed of larger condensates in sizes than ones in the first population (orange, **Figure 4D**). On the other hand, the size distributions of condensates formed from rigid filaments were unimodal with one population whose size was close to the size that could be formed by four filaments themselves (blue, **Figure 4D**). This may be because the geometry of rigid filaments provides limited accessibility to freely diffusing PFKL tetramers while elastic filaments may be evolved to either of two populations due to their flexibility. Second, when the enzyme-enzyme interaction was set as *ε* = 4.5 × 10^−23^ (red and green, **Figure 4D**), regardless of the flexibility of computed filaments, subpopulations whose sizes were greater than 400 were distinctively formed. As the length of filaments got longer, the width of the subpopulation’s distribution containing large condensates (i.e., > 400 in size) became narrower while the population displaying smaller condensates (i.e., < 400 in size) was increased. Therefore, the observed trend at *ε* = 4.5 × 10^−23^ was similar in both elastic and rigid filaments.

Collectively, we identified *in silico* several biologically relevant parameters of PFKL (i.e., its intermolecular interaction strength, its effective concentration, its multivalency, and its pre-organization prior to condensate formation) that would regulate the formation of PFKL condensates and their sizes in cells. Accordingly, we propose that, when the scaffolder enzyme of a multienzyme glucosome condensate, PFKL, is subcellularly controlled to present relatively a stronger interaction or a higher effective concentration, regulated to be located in a higher multivalency mimicking environment, or organized to form filamentous or similar structures in a cell, the formation of glucosome condensates would be promoted to orchestrate glucose flux in conjunction with other cellular processes in human cells (Jeon *et al*., 2022a; Jeon *et al*., 2018; Jeon *et al*., 2022b; Kennedy *et al*., 2022; Kohnhorst *et al*., 2017). Considering our study was done *in silico* with stochastic modeling approaches, we offer a possibility that any metabolic enzymes might be able to form spatially resolved condensates, but at different degrees, in human cells particularly when their biological and physical properties are perturbed acutely and/or dysregulated chronically in human cells.

## Discussion

In this work, we developed a stochastic model to describe how PFKL is organized to initiate the formation of a multienzyme metabolic condensate, the glucosome. The model was constructed based on the Langevin dynamics, in which each PFKL tetramer was governed by forces originated from its random diffusion, friction, and interaction. In this model, the shape of the PFKL tetramer was assumed as a point particle and the dynamics of PFKL filaments was dictated by a bead-spring model (Geyer, 2011). Accordingly, we disclose the importance of enzyme-specific properties of PFKL and their tunability in formation of PFKL condensates as an initiation step of glucosome formation.

Previously, we demonstrated the intracellular formation of glucosome condensates was governed by LLPS in living cells (Kennedy *et al*., 2022; Kohnhorst *et al*., 2017) similarly as other biomolecular condensates do (Banani *et al*., 2017). However, due to the nature of our findings from live cells, it has been challenging to characterize which parameters would contribute to phase separation of PFKL and thus formation of glucosome condensates. In this work, we took advantage of *in silico* approaches and investigated the impacts of enzyme-specific biological properties of PFKL on the initiation step of glucosome formation. We then found that various parameters showed a threshold and a defined range at which condensates formation and their size distributions were regulated. We also noticed that both freely diffusing PFKLs and bound PFKLs to condensates coexisted during the simulation of our stochastic model. This is indeed consistent with what we have observed in living human cells (Kennedy *et al*., 2022; Kohnhorst *et al*., 2017), where two populations of PFKLs (i.e., inside and outside a condensate) often coexist depending on subcellular environment or available exogenous stimuli where enzymatic properties of PFKL are regulated. Collectively, we demonstrate that the biological properties of PFKL which would be intrinsically under controls by various cellular processes, including but not limited to signaling pathways (Jeon *et al*., 2022a) and a cell cycle (Jeon *et al*., 2022b), are critical determinants at the initiation step for glucosome formation in human cells.

We also demonstrate here that a pre-organization of PFKL into a filamentous structure can overcome various limitations in order for PFKL to form glucosome condensates. As demonstrated in **Figures 1 and 4A**, when PFKLs were organized into filaments, the probability of PFKL being formed into a condensate was significantly increased from nearly 0 % to > 97% at *ε* = 4.5 × 10^−23^. Even, when PFKL had no chance to be organized into a condensate at *ε* = 3.3 × 10^−23^, the pre-organization of PFKL into filaments significantly enhanced the probability of PFKL being formed into a condensate (i.e., 0% to > 65% with elastic filaments containing at least 4 PFKL tetramers or with rigid filaments having at least 5 PFKL tetramers, respectively) (**Figure 4A**). Since many metabolic enzymes now have been discovered to display various forms of mesoscale subcellular structures (e.g., filaments, rods, rings, condensates, etc) (Wilson and Gitai, 2013), a pre-organization of a given enzyme may be formed as a substructure for such mesoscale structures in cells. Collectively, it appears that an initial pre-organization of an enzyme appears to be the most critical determinant for condensate formation in a cell.

It is also important to emphasize here that we have analyzed contributions of LLPS-associated parameters of PFKL to size distribution of its condensates. Although the importance of biomolecular condensates’ sizes is generally not well understood at this time, glucosome condensates have been demonstrated to control glucose flux in a condensate size-dependent manner (Jeon *et al*., 2018; Kennedy *et al*., 2022; Kohnhorst *et al*., 2017). Briefly, small-sized glucosomes (< 0.1 μm^2^) defined under fluorescence live-cell imaging are responsible for operating glycolysis for energy metabolism (Jeon *et al*., 2018; Kennedy *et al*., 2022) while medium-and large-sized glucosomes (0.1 - 3 μm^2^ and > 3 μm^2^, respectively) shunt glucose flux into building block biosynthesis (i.e., the pentose phosphate pathway and serine biosynthesis) (Jeon *et al*., 2018; Kohnhorst *et al*., 2017). Now, we showed, for instance, that the number of condensates having > 400 PFKLs was increased while the intermolecular interaction strength (*ε*) was increased (**Figure 1B**), suggesting the promotion of glucosome activities into building block biosynthesis. In contrast, when the intermolecular interaction strength was very weak like *ε* = 3.3 × 10^−23^ (**Figure 4D**), all the condensates had < 100 PFKL tetramers (i.e., small-sized glucosomes), thereby driving glucose flux into energy metabolism. Meanwhile, at the threshold of the intermolecular interaction strength (i.e., *ε* = 4.5 × 10^−23^, **Figure 1A**), additional parameters appear to influence condensate sizes and thus functions. Collectively, we propose that the biological parameters of PFKL at a given time and subcellular location (i.e., its intermolecular interaction strength, its effective concentration, its multivalency, and its pre-organization) are not only important for condensates formation but also their functional contributions to cellular metabolism in human cells.

Nevertheless, we understand that our current model needs to be improved to advance our understanding of reversible dynamics of glucosomes in a cell. For example, our model does not include a dissociation event of a condensate yet. Accordingly, once a PFKL condensate is formed, the condensate would be programed to keep growing as time goes on due to our application of the interaction potential that uses a distance-dependent attractive or repulsive force. Also, our model does not include other enzymes that are known to be involved in glucosome formation such as fructose 1,6-bisophosphatase, pyruvate kinase muscle-type 2, and phosphoenolpyruvate carboxykinase 1 (Kohnhorst *et al*., 2017). Nevertheless, our current model is particularly advantageous to explain how PFKL initiates the formation of a condensate and thus describe an initiation step of glucosome formation. Therefore, we envision that our model would be advanced to explain a next stage of glucosome formation, namely its dynamic maturation step, by including both fusion and fission events of condensates with additional enzyme components.

## Acknowledgments

This work is supported by the National Institutes of Health: R01GM125981 (SA), R03CA219609 (SA), R01GM134086 (MK) and the National Science Foundation: DMS-1620403 (H-W.K.). The content is solely the responsibility of the authors and does not necessarily represent the official views of the National Institutes of Health.

## Author contributions

H-W.K. and L.N. performed numerical simulations, M.K. carried out statistical analysis, H-W.K., S.A., and M.K. conceived the project, analyzed, and discussed the results, and wrote the manuscript.

## Data availability

All data are available within the main manuscript.

